# A Unified Framework for Systematic Curation and Evaluation of Aging Biomarkers

**DOI:** 10.1101/2023.12.02.569722

**Authors:** Kejun Ying, Seth Paulson, Alec Eames, Alexander Tyshkovskiy, Siyuan Li, Martin Perez-Guevara, Mehrnoosh Emamifar, Maximiliano Casas Martínez, Dayoon Kwon, Anna Kosheleva, Michael P. Snyder, Dane Gobel, Chiara Herzog, Jesse R. Poganik, Biomarker of Aging Consortium, Mahdi Moqri, Vadim N. Gladyshev

## Abstract

Aging biomarkers are essential for understanding and quantifying the aging process and developing targeted longevity interventions. However, validation of these tools has been hindered by the lack of standardized approaches for cross-population validation, disparate biomarker designs, and inconsistencies in dataset structures. To address these challenges, we developed Biolearn, an open-source library that provides a unified framework for the curation, harmonization, and systematic evaluation of aging biomarkers. Leveraging Biolearn, we conducted a comprehensive evaluation of various aging biomarkers across multiple datasets. Our systematic approach involved three key steps: (1) harmonizing existing and novel aging biomarkers in standardized formats; (2) unifying public datasets to ensure coherent structuring and formatting; and (3) applying computational methodologies to assess the harmonized biomarkers against the unified datasets. This evaluation yielded valuable insights into the performance, robustness, and generalizability of aging biomarkers across different populations and datasets. The Biolearn python library, which forms the foundation of this systematic evaluation, is freely available at https://Bio-Learn.github.io. Our work establishes a unified framework for the curation and evaluation of aging biomarkers, paving the way for more efficient and effective clinical validation and application in the field of longevity research.

## Introduction

Development and validation of robust biomarkers of aging (BoAs) have become key focal points in aging research, driven by the growing recognition of aging as a fundamental driver of chronic diseases and mortality. Numerous biomarkers have been proposed to quantify biological age and elucidate the biological processes underlying aging. However, clinical validation of BoAs remains a significant challenge due to heterogeneity in their formulations and disparate structures of validation datasets across populations ^1,2^.

Since the introduction of composite omic biomarkers of aging, exemplified by Horvath’s pioneering work on DNA methylation aging clocks ^3^, the field has witnessed a rapid expansion in the repertoire of aging biomarkers. These biomarkers now span a wide array of omic modalities, including epigenomics, transcriptomics, and proteomics ^4–9^. Omic biomarkers provide a compre-hensive view of the molecular changes associated with aging, offering valuable insights into the aging process and its impact on human health.

Among the various classes of omic biomarkers, DNA methylation-based clocks are currently the most advanced and robust tools for estimating biological age. These human clocks, such as the Horvath multi-tissue clock, DunedinPACE ^5^, GrimAge ^10^, PhenoAge ^11^, causality-enriched DamAge/AdaptAge ^12^, and the PRC2 clock ^13^, have demonstrated significant associations with age-related conditions and mortality, highlighting the intricate relationship between epigenetic modifications and aging trajectories ^6,10,14,15^. However, the diverse formulations of these biomarkers and inconsistencies in dataset structures across different populations pose substantial challenges for their systematic cross-population validation and benchmarking, which are crucial steps toward their clinical translation.

Publicly available datasets, such as those from the Gene Expression Omnibus (GEO) ^16^, the National Health and Nutrition Examination Survey (NHANES), and the Framingham Heart Study (FHS), hold immense potential for accelerating the validation of BoAs. However, the lack of a standardized framework that can accommodate the heterogeneous nature of these datasets hinders their effective utilization for this purpose. There is a pressing need for a unified platform that can seamlessly integrate and analyze various BoAs across datasets with harmonized structures. Such a platform would revolutionize the validation process, facilitate the discovery of novel biomarkers, and provide a structured avenue for community-driven efforts in advancing the field of aging biology.

To address this need, we developed Biolearn, an open-source Python library that provides a unified framework for the curation, harmonization, and systematic evaluation of aging biomarkers (Figure 1a). Biolearn supports biomarkers based on multiple different biological data modalities and serves as an innovative tool that harmonizes existing BoAs, structures and formats human datasets and offers computational methodologies for assessing biomarkers against these datasets. By enabling the integration and analysis of diverse BoAs and datasets, Biolearn aims to accelerate the development and validation of BoAs, fostering a community-driven approach to aging research.

**Figure 1.**
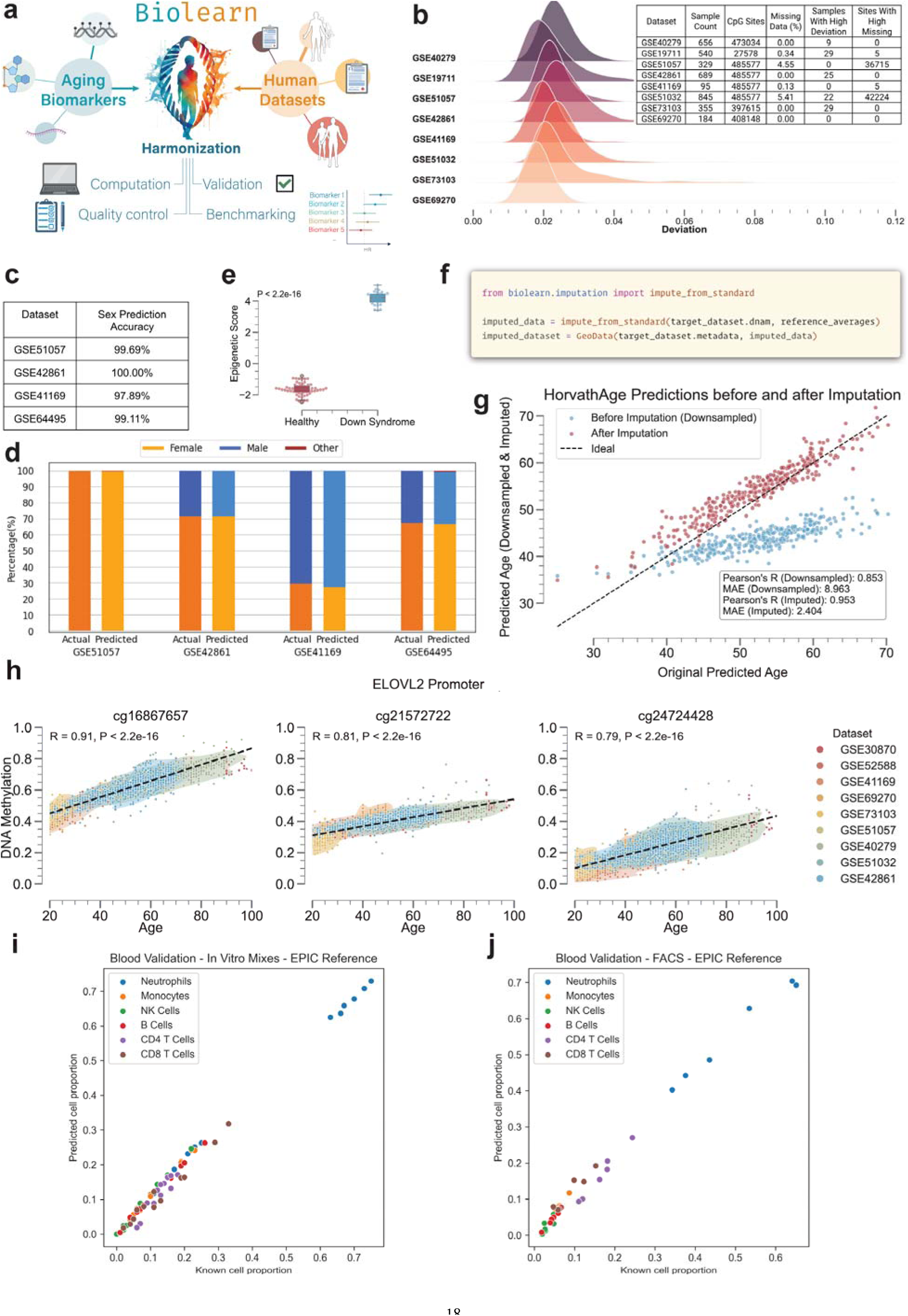
Biolearn: A harmonization framework for aging biomarkers and biological datasets. **a.** Overview of Biolearn’s functionalities, including biomarker discovery, dataset harmonization, and imputation. **b.** Quality control results are visualized as a ridge density plot showing the distribution of sample deviations from the mean of the population. The accompanying table lists key metadata for each dataset, including sample count, age statistics, and disease status. **c.** Comparison of actual vs. predicted sex distributions in four datasets (GSE51057, GSE42861, GSE41169, and GSE64495), with prediction accuracies listed in the accompanying table. **d.** Stacked bar graphs displaying the actual and predicted sex distributions for each dataset, with female, male and other categories shown. **e.** Epigenetic score distribution of healthy individuals compared to those with Down Syndrome (P < 2.2e-16, Wilcoxon rank-sum test). **f.** Python code snippet illustrating the usage of Biolearn to import, preprocess and impute missing data in a target dataset using a reference dataset. **g.** Scatter plot showing a significant inverse correlation between the Horvath epigenetic age predictions and imputed Hannum age predictions across multiple datasets (Pearson’s R = 0.999, MAE = 0.325). **h.** DNA methylation levels of three age-associated CpG sites (cg16867657, cg21572722, cg24724428) plotted against chronological age for 9 datasets, with points colored by dataset. Robust linear regressions are shown, with the correlation coefficient (R) and p-value displayed for each CpG site. **i,j.** Deconvolution results for the DeconvoluteBloodEPIC method applied to a dataset with known cell proportions from in vitro mixes (**i**) and with FACS (**j**). The estimated cell type proportions are compared to the true proportions.

## Results

### Harmonization of Biomarkers of Aging and Datasets

We harmonized a comprehensive set of 39 well-established epigenetic, transcriptomic, and clinical biomarkers (Table 1) and implemented these BoAs in Biolearn, representing the largest collection of BoAs in a single package to date. We have validated the implementation of these biomarkers with their respective developers to ensure accuracy and reliability. The epigenetic biomarkers encompass a wide range of categories, including: (1) Chronological clocks: Horvath’s multi-tissue clock and Hannum’s blood clock ^3,17^; (2) Healthspan and mortality-related clocks: GrimAge, GrimAge2, PhenoAge, and Zhang clock ^10,11,18,19^; (3) Biomarkers of the rate of aging: DunedinPoAm38 and DunedinPACE ^5,20^; (4) Causality-enriched clocks: Ying’s CausAge, DamAge, and AdaptAge ^12^; and (5) Various other clocks, including DNAm-based biomarkers and disease predictors, transcriptomic clocks, and clinical clocks (Table 1).

**Table 1.** Harmonized biomarkers in Biolearn.

| Biomarker | Year | Tissue | Predicts | Omic Type |
| --- | --- | --- | --- | --- |
| HorvathV1 <sup>3</sup> | 2013 | Multi-tissue | Age (Years) | DNA Methylation |
| Hannum <sup>17</sup> | 2013 | Blood | Age (Years) | DNA Methylation |
| Lin <sup>39</sup> | 2016 | Blood | Age (Years) | DNA Methylation |
| PhenoAge <sup>11</sup> | 2018 | Blood | Age (Years) | DNA Methylation |
| HorvathV2 <sup>40</sup> | 2018 | Skin, blood | Age (Years) | DNA Methylation |
| PEDBE <sup>41</sup> | 2019 | Buccal | Age (Years) | DNA Methylation |
| Zhang_10 <sup>19</sup> | 2019 | Blood | Mortality Risk | DNA Methylation |
| GrimAge <sup>10</sup> | 2019 | Blood | Age Adjusted by Mortality Risk (Years) | DNA Methylation |
| GrimAge2 <sup>18</sup> | 2022 | Blood | Age Adjusted by Mortality Risk (Years) | DNA Methylation |
| DunedinPoAm38 <sup>20</sup> | 2020 | Blood | Aging Rate (Years/Year) | DNA Methylation |
| DunedinPACE <sup>5</sup> | 2022 | Blood | Aging Rate (Years/Year) | DNA Methylation |
| DNAmTL <sup>42</sup> | 2019 | Blood, Adipose | Telomere Length | DNA Methylation |
| Knight <sup>43</sup> | 2016 | Cord Blood | Gestational Age | DNA Methylation |
| LeeControl <sup>44</sup> | 2019 | Placenta | Gestational Age | DNA Methylation |
| LeeRefinedRobust <sup>44</sup> | 2019 | Placenta | Gestational Age | DNA Methylation |
| LeeRobust <sup>44</sup> | 2019 | Placenta | Gestational Age | DNA Methylation |
| YingCausAge <sup>12</sup> | 2022 | Blood | Age (Years) | DNA Methylation |
| YingDamAge <sup>12</sup> | 2022 | Blood | Age (Years) | DNA Methylation |
| YingAdaptAge <sup>12</sup> | 2022 | Blood | Age (Years) | DNA Methylation |
| SmokingMcCartney <sup>45</sup> | 2018 | Blood | Smoking Status | DNA Methylation |
| AlcoholMcCartney <sup>45</sup> | 2018 | Blood | Alcohol Consumption | DNA Methylation |
| BMI_McCartney <sup>45</sup> | 2018 | Blood | BMI | DNA Methylation |
| EducationMcCartney <sup>45</sup> | 2018 | Blood | Educational Attainment | DNA Methylation |
| TotalCholesterolMcCartney <sup>45</sup> | 2018 | Blood | Total Cholesterol | DNA Methylation |
| HDLCholesterolMcCartney <sup>45</sup> | 2018 | Blood | HDL Cholesterol | DNA Methylation |
| LDLCholesterolMcCartney <sup>45</sup> | 2018 | Blood | LDL with Remnant Cholesterol | DNA Methylation |
| BodyFatMcCartney <sup>45</sup> | 2018 | Blood | Percentage Body Fat | DNA Methylation |
| BMI_Reed <sup>46</sup> | 2020 | Blood | BMI | DNA Methylation |
| ProstateCancerKirby <sup>47</sup> | 2017 | Prostate | Prostate Cancer Status | DNA Methylation |
| HepatoXu <sup>48</sup> | 2017 | Circulating DNA | Hepatocellular Carcinoma Status | DNA Methylation |
| CVD_Westerman <sup>49</sup> | 2020 | Blood | Coronary Heart Disease Status | DNA Methylation |
| AD_Bahado-Singh <sup>50</sup> | 2021 | Blood | Alzheimer's Disease Status | DNA Methylation |
| DepressionBarbu <sup>51</sup> | 2021 | Blood | Depression Risk | DNA Methylation |
| Phenotypic Age <sup>11</sup> | 2018 | Blood | Phenotypic Age | Clinical Biomarker |
| Mahalanobis Distance <sup>34</sup> | 2023 | Blood | Mahalanobis Distance | Clinical Biomarker |
| Peters_tAge <sup>30</sup> | 2015 | Blood | Age (Years) | RNA |
| Multispecies-blood_tAge | 2024 | Blood | Relative Age (Years) | RNA |
| Human-blood_tAge | 2024 | Blood | Relative Age (Years) | RNA |
| Human-multitissue_tAge | 2024 | Multi-tissue | Relative Age (Years) | RNA |

The harmonization of these diverse biomarkers is crucial for enabling consistent and reproducible analyses across different datasets, facilitating cross-population validation studies, and advancing our understanding of the aging process. To achieve this, all biomarkers were formatted into standardized input structures, ensuring their consistent application across disparate datasets. The harmonization process involved collecting and unifying the annotation of clock specifications, such as tissue type, predicted age range, and source references. This meticulous approach guarantees transparent and reproducible analyses, enabling researchers to readily compare and interpret results across different studies and populations.

To further support ongoing research in this field, we developed and implemented an open-source framework. This framework provides a standardized format for epigenetic biomarkers, facilitating the seamless integration and comparison of any future aging clocks and biomarkers that are developed. By providing a unified platform for the harmonization and analysis of aging biomarkers, this framework aims to foster collaboration and innovation. Researchers can easily contribute new biomarkers, compare their performance against existing ones, and explore their potential applications in various datasets. This collaborative approach is essential for accelerating progress in the field and developing more accurate and robust biomarkers of aging.

To facilitate cross-population validation studies using publicly available data, we harnessed Biolearn’s capabilities to integrate and structure multiple public datasets (Table 2). The structured datasets were refined to enable a shared analysis platform, addressing the challenges of data heterogeneity and formatting inconsistencies ^21,22^. With this capacity, Biolearn is used as the backend of ClockBase for epigenetic age computation ^22^, enabling the systemic harmonization of over 200,000 human samples from Gene Expression Omnibus (GEO) array data.

**Table 2.** Harmonized datasets in Biolearn.

| ID | Title | Format | Samples | Age Present | Sex Present |
| --- | --- | --- | --- | --- | --- |
| GSE40279 | Genome-wide Methylation Profiles Reveal Quantitative Views o... | Illumina450k | 656 | Yes | Yes |
| GSE19711 | Genome wide DNA methylation profiling of United Kingdom Ovar... | Illumina27k | 540 | Yes | No |
| GSE51057 | Methylome Analysis and Epigenetic Changes Associated with Me... | Illumina450k | 329 | Yes | Yes |
| GSE42861 | Differential DNA methylation in Rheumatoid arthritis | Illumina450k | 689 | Yes | Yes |
| GSE41169 | Blood DNA methylation profiles in a Dutch population | Illumina450k | 95 | Yes | Yes |
| GSE51032 | EPIC-Italy at HuGeF | Illumina450k | 845 | Yes | No |
| GSE73103 | Many obesity-associated SNPs strongly associate with DNA met... | Illumina450k | 355 | Yes | Yes |
| GSE69270 | Aging-associated DNA methylation changes in middle-aged indi... | Illumina450k | 184 | Yes | No |
| GSE36054 | Methylation Profiling of Blood DNA from Healthy Children | Illumina450k | 192 | No | No |
| GSE64495 | DNA methylation profiles of human blood samples from a sev-er... | Illumina450k | 113 | Yes | Yes |
| GSE30870 | DNA methylomes of Newborns and Nonagenarians | Illumina450k | 40 | Yes | No |
| GSE52588 | Identification of a DNA methylation signature in blood from ... | Illumina450k | 87 | Yes | Yes |
| GSE157131 | Methylation data from stored peripheral blood leukocytes fro... | IlluminaEPIC | 946 | Yes | Yes |
| GSE132203 | DNA Methylation (EPIC) from the Grady Trauma Project | IlluminaEPIC | 795 | Yes | Yes |
| GSE134080 | RNASeq whole blood of Dutch 500FG cohort | IlluminaHiSeq2500 | 100 | Yes | Yes |
| NHANES | National Health and Nutrition Examination Survey | Phenotypic | 2877 | Yes | Yes |
| FHS | Framingham Heart Study | Phenotypic | 4434 | Yes | Yes |

### Quality Control, Imputation, and Deconvolution

Biolearn provides a comprehensive toolkit for data preprocessing, normalization, and cell-type deconvolution (Figure 1a). Quality control metrics, such as sample deviation from the population mean, missingness, and the number of sites with a high percentage of missingness, can be readily visualized (Figure 1b). This functionality enables researchers to identify potential outliers or problematic samples and make informed decisions about data inclusion and exclusion criteria.

Moreover, Biolearn enables the prediction of sample sex from DNA methylation data with high accuracy, which can be compared against actual sex distributions (Figure 1c,d)^23^. This feature is particularly useful for identifying potential sample mislabeling or investigating sex-specific effects in aging research. We also implemented predictors of common traits, including smoking, BMI, and epigenetic scores for diseases like Down Syndrome (Figure 1e)^24^. These predictors allow researchers to explore the associations between aging biomarkers and various lifestyle factors or disease conditions, providing valuable insights into the complex interplay between aging and health.

Missing DNA methylation data can be easily imputed with different methods in just a few lines of code (Figure 1f,g) ^25^, ensuring that researchers can make the most of the available data and minimize the impact of missing values on their analyses. Biolearn also facilitates the integration of multiple datasets for large-scale analyses, such as the comparison of DNA methylation levels of CpG sites across different datasets (Figure 1h). This feature allows researchers to investigate the consistency and reproducibility of aging biomarkers across diverse populations and experimental settings, strengthening the robustness and generalizability of their findings.

In addition to these features, Biolearn offers a deconvolution tool that estimates the proportion of cell types in a given sample based on a single bulk-level methylation measurement. Biolearn provides two modes for deconvolution, optimized for the 450K (DeconvoluteBlood450K) and EPIC (DeconvoluteBloodEPIC) methylation platforms, respectively ^26^. These modes are designed for estimating cell proportions in blood methylation samples and account for technology-specific biases that can affect the accuracy of deconvolution ^27^. The reference methylation matrices for each mode consist of methylation profiles for six cell types representing the most abundant cell types found in the blood: neutrophils, monocytes, natural killer cells, B cells, CD4+ T cells, and CD8+ T cells ^28,29^.

We benchmarked the accuracy of our deconvolution tool using datasets with known cell proportions assessed via fluorescence-activated cell sorting (FACS) and *in vitro* cell mixing ^29^. Both deconvolution methods generated accurate predictions that matched known cell proportions (Figure 1i,j). These results demonstrate the reliability and utility of Biolearn’s deconvolution feature, which can help researchers account for cellular heterogeneity in their analyses and gain insights into the cell type-specific contributions to aging biomarkers.

### Systematic Evaluation of the Biomarkers of Aging

To demonstrate the utility of Biolearn in facilitating the systematic evaluation of aging biomarkers, we conducted a comprehensive benchmarking analysis of various epigenetic aging clocks across multiple datasets. By leveraging Biolearn’s harmonized dataset library and standardized clock implementations, we assessed the performance, robustness, and generalizability of these clocks in diverse biological contexts and populations (Figure 2). Our analysis included a wide range of datasets, spanning the Human Aging Rates Study (GSE40279, N = 656), EPIC-Italy (GSE51032, N = 845), RA Case-control Cohort (GSE42861, N = 689), Dutch Schizophrenia Case-control Cohort (GSE41169, N = 95), Obesity Genetics Study (GSE73103, N = 355), Developmental Disorder Study (GSE64495, N = 113), African American GENOA (GSE157131, N = 1218), and Grady Trauma Project (GSE132203, N = 795) (Figure 2a). These diverse datasets allowed us to evaluate the performance of the biomarkers across various age ranges, ethnicities, and disease states. Biolearn’s user-friendly interface and efficient data handling capabilities streamlined data loading and implementation of aging clocks (Figure 2b). This highlights the library’s potential to accelerate the development and validation of novel aging biomarkers by providing a standardized framework for their evaluation.

**Figure 2.**
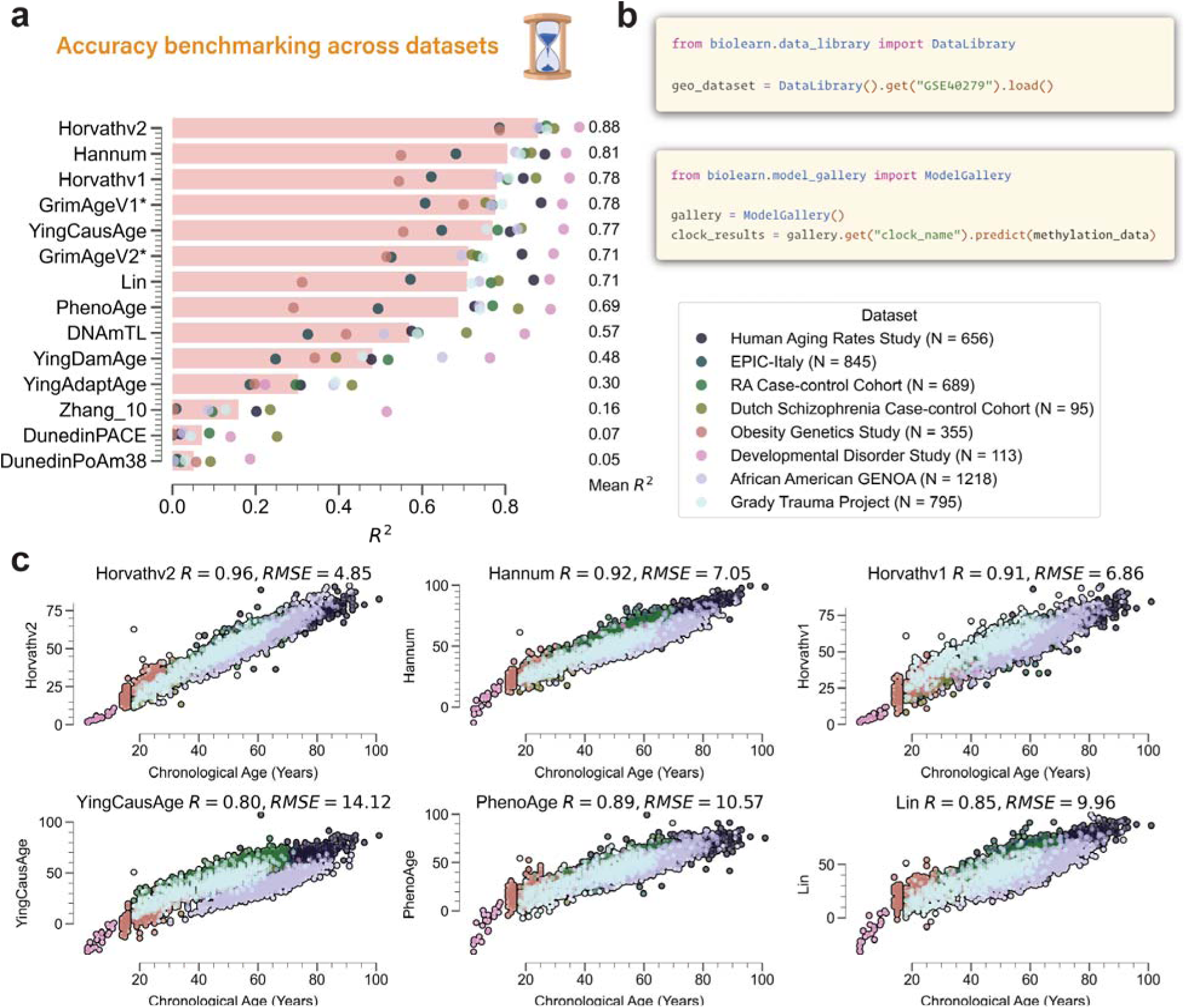
Systematic evaluation of epigenetic aging clocks across diverse datasets using Biolearn. **a.** Benchmarking of aging clocks across eight datasets, with mean R^2^ values indicating the average performance of each clock. The dots colored based on the dataset represent the R^2^ measured in each individual dataset. The datasets span various ethnicities and populations, enabling a comprehensive assessment of clock robustness and generalizability. **b.** Code snippets demonstrating the streamlined curated data loading and implementation of aging clocks using the Biolearn library, highlighting its user-friendly interface and efficient data handling. **c.** Scatter plots depicting the relationship between predicted epigenetic age and chronological age for six representative clocks across different datasets represented by different colors. The plots showcase the performance of each clock, with Pearson’s R and RMSE provided. The datasets cover a wide age range, allowing for a thorough evaluation of clock accuracy and applicability across diverse age groups and biological contexts. The datasets shown include the Human Aging Rates Study (GSE40279), UKOPS (GSE19711), Dutch Schizophrenia Case-control Cohort (GSE41169), Obesity Genetics Study (GSE73103), Young Finns Study (GSE69270), and Newborns and Nonagenarians Study (GSE30870).

The benchmarking results (Figure 2a-c) revealed that the HorvathV2 clock (i.e., skin and blood clock) exhibited the highest overall accuracy in terms of predicting chronological age, with a mean R^2^ of 0.88 across all datasets, followed closely by the Hannum clock (R^2^ = 0.81), the Horvath1 clock (R^2^ = 0.78) and YingCausAge clock (R^2^ = 0.77). These findings suggest that these four clocks are the most robust and generalizable across diverse biological contexts and agerelated conditions. Note that the GrimAgeV1 and GrimAgeV2 clocks use the age of the sample as the predictor, therefore they cannot be compared directly to the other clocks. These results highlight the applicability of these clocks across diverse age ranges and biological contexts, further emphasizing the importance of systematic evaluation in identifying the most suitable biomarkers for specific research questions or clinical applications.

Overall, our findings demonstrate the value of Biolearn in enabling the systematic evaluation of epigenetic aging clocks across multiple datasets. By providing a standardized framework for clock implementation and evaluation, Biolearn facilitates the identification of robust and generalizable aging biomarkers, paving the way for their translation into clinical settings and advancing our understanding of the aging process.

### Mortality and Morbidity Risk Analysis

It is also important to evaluate the predictive power of epigenetic aging biomarkers in predicting aging-associated outcomes such as mortality risk. Here, we conducted the most comprehensive evaluation of 17 representative epigenetic clock models to the Normative Aging Study (NAS) dataset (N = 1,488, 38.8% deceased), comparing their performance in predicting mortality risk (Figure 3). The biomarkers showed a strong correlation with chronological age in both NAS (Figure 3a).

**Figure 3.**
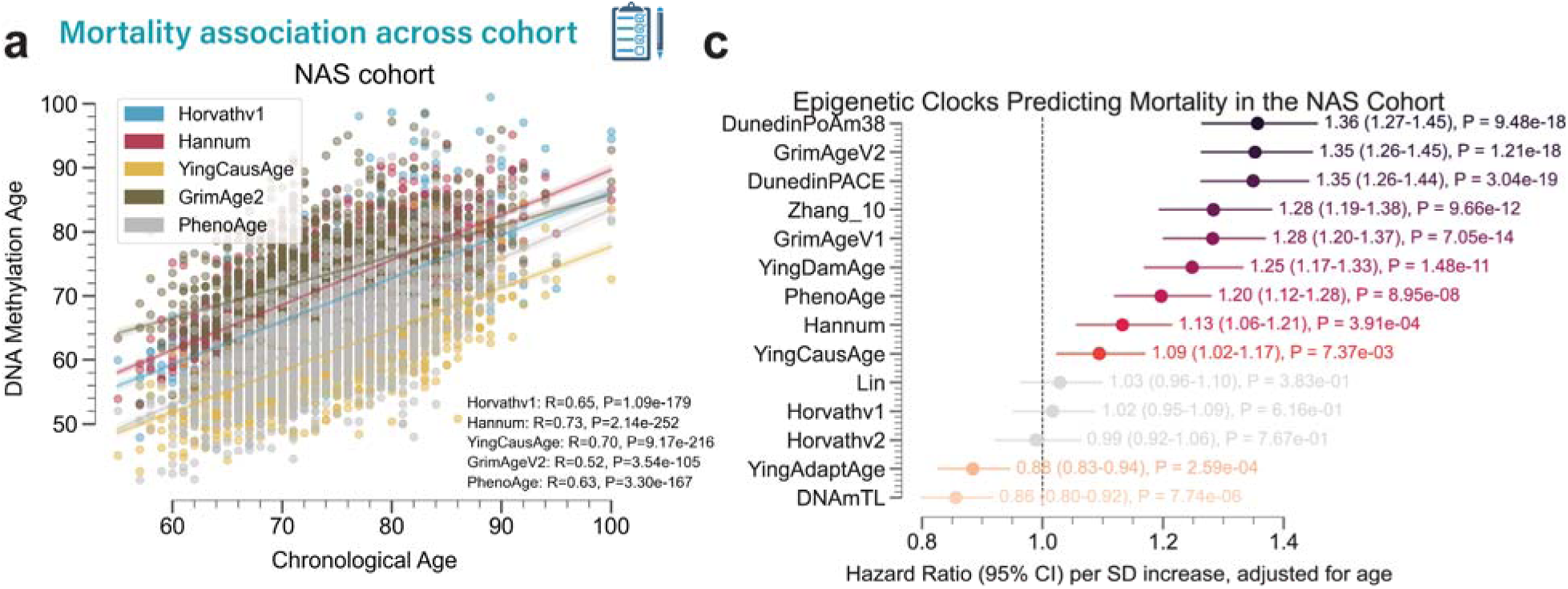
Comprehensive analysis of epigenetic clocks in predicting mortality. **a-b.** Scatter plots illustrate the relationship between DNA methylation age and chronological age for the NAS cohort (N = 1,488, 38.8% deceased, **a**). The five most representative clock models are shown for each cohort. The accuracy of each clock is indicated by Pearson’s R value, with associated P-values based on two-sided tests. **c-d.** Forest plot presenting the hazard ratios (HR) of Cox proportional hazards regression models predicting mortality risk for the NAS cohort (**c**) as predicted by 14 epigenetic clock models, adjusted for age. The hazard ratios are shown per standard deviation increase of the biomarker, with 95% confidence intervals displayed as error bars. P-values are based on two-sided tests. The insignificant results (P > 0.05) are colored gray. **e.** Scatter plot comparing the Pearson’s correlation coefficient with chronological age (x-axis) and the hazard ratio of mortality risk (y-axis) for each aging clock in NAS. The points are colored based on the cohorts. The regression line and 95% confidence intervals are shown. Pearson’s R-value and P-value are shown at the top of the plot.

We then examined their performance in predicting mortality risk using Cox Proportional Hazards analysis, adjusted for age and sex (Figure 3c, d). In the NAS cohort (Figure 3c), the top-performing clock in terms of hazard ratio (per standard deviation increase of the biomarker) was DunedinPoAm38 (HR = 1.38, P = 9.48e-18), followed closely by GrimAgeV2 (HR = 1.35, P = 1.21e-18) and DunedinPACE (HR = 1.35, P = 3.04e-19). Other clocks with strong predictive value include Zhang_10 (HR = 1.28, P = 9.66e-12), GrimAgeV1 (HR = 1.29, P = 7.05e-14), YingDamAge (HR = 1.25, P = 1.48e-11), PhenoAge (HR = 1.20, P = 8.96e-08), Hannum (HR = 1.13, P = 3.91e-04), and YingCausAge (HR = 1.09, P = 7.37e-03).

We further assessed the association between the predictive power of the epigenetic clocks for chronological age and mortality risk. Interestingly, in both cohorts, we observed a negative but insignificant correlation between Pearson’s R with chronological age and hazard ratio of mortality risk (Figure 3e). This suggests that the predictive power of epigenetic clocks on mortality risk, after adjusting for age and sex, is independent of their ability to predict chronological age, highlighting the importance of interpreting the meaning of age deviation (AgeDev) with caution for aging biomarkers. We also observed strong heterogeneous associations of epigenetic clocks with mortality risk in different cohorts. The analysis demonstrates the utility of Biolearn in facilitating systematic evaluation of epigenetic clocks in predicting mortality across multiple cohorts, emphasizing the importance of systematic evaluation in identifying the most suitable biomarkers for specific applications.

Besides mortality, it is also important to evaluate aging biomarkers in predicting various other clinically relevant aging outcomes. We analyzed the associations between 14 aging biomarkers and six event categories (Stroke, Dementia, Operation, Lifespan, Cancer, and Healthspan, which is defined by the first incidence of any event) in the NAS cohort (Figure 4a). To assess the predictive power of these biomarkers, we performed Cox Proportional Hazards analyses, adjusted for age, and calculated the hazard ratios (HR) per standard deviation increase for each biomarker (Figure 4b-f).

**Figure 4.**
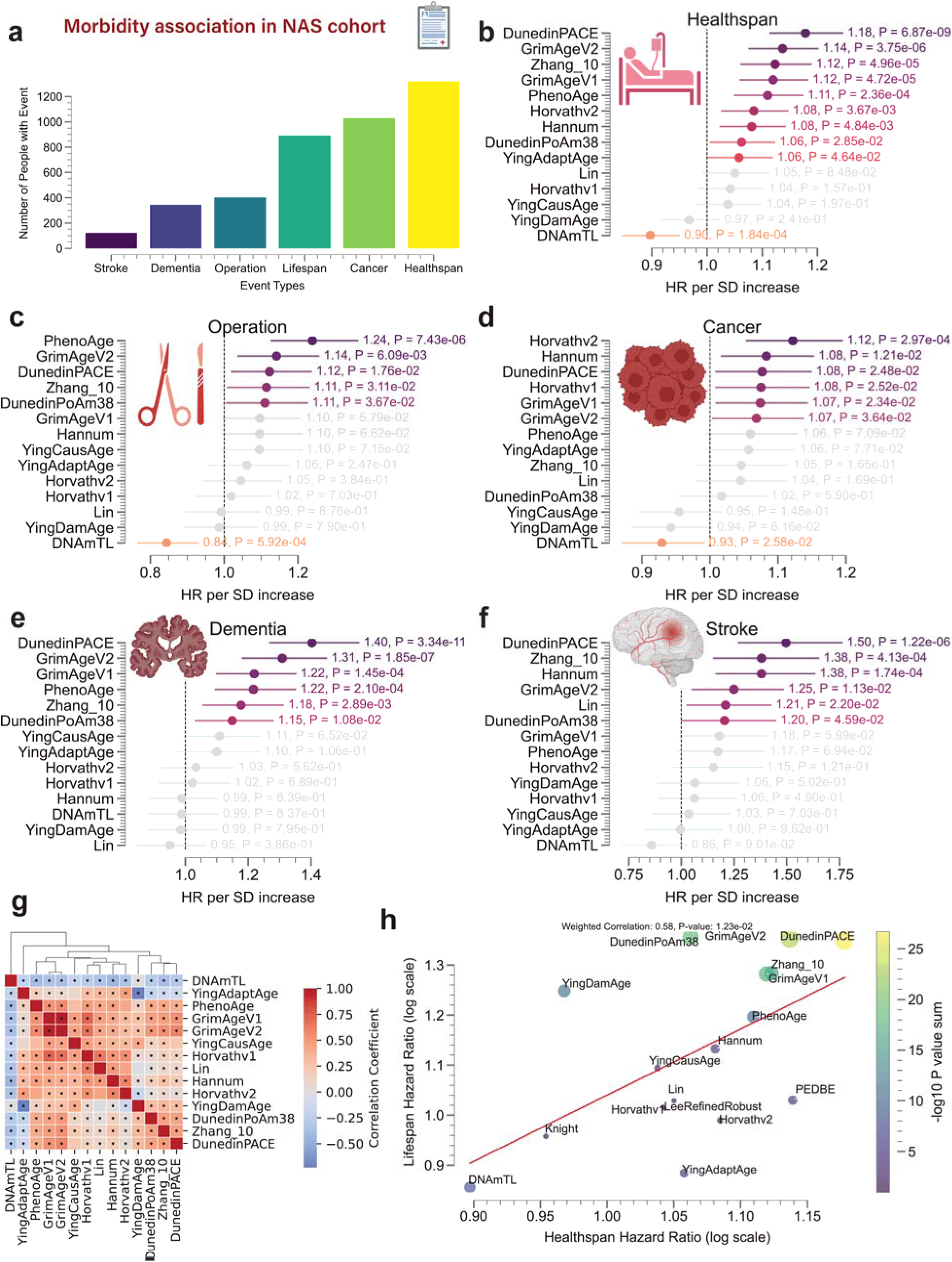
Comprehensive analysis of aging biomarkers in predicting clinical outcomes. **a.** Bar plot displaying the number of people with events across six categories (Stroke, Dementia, Operation, Lifespan, Cancer, and Healthspan) in the NAS cohort. **b-f.** Forest plots presenting the hazard ratios (HR) per standard deviation increase for various aging biomarkers in predicting specific clinical outcomes: Healthspan (b), Operation (c), Cancer (d), Dementia (e), Stroke (f), and individual event types (g). The 95% confidence intervals are displayed as error bars, and P-values are based on two-sided tests. **g.** Correlation matrix showcasing the associations between different aging biomarkers in the NAS cohort, after adjusting for age. The color intensity represents the strength of the correlation, with red indicating positive correlations and blue indicating negative correlations. The significant pairs are annotated with black dots. **h.** Scatter plot comparing the Healthspan hazard ratios (log scale) against the log2-transformed lifespan hazard ratios for various aging biomarkers. The plot reveals the relationship between the predictive power of biomarkers for healthspan and lifespan. The size of the points represents the significance of the P-values, with larger points indicating higher significance. The inverse variance weighted correlation and P-values are shown.

Across five clinical outcomes tested, DunedinPACE was the strongest predictor for three of the outcomes, namely healthspan (HR = 1.18), dementia (HR = 1.40), and stroke (HR = 1.50). PhenoAge was the strongest predictor for surgery (HR = 1.24), and HorvathV2 was the strongest predictor for cancer (HR = 1.12). These results suggest considerable heterogeneity of aging biomarkers in predicting different clinical outcomes.

We further investigated the associations between the AgeDev term of aging biomarkers after adjusting for age in the NAS cohort and found strong positive correlations among most epigenetic clocks (Figure 4g). We observed two main clusters: (1) PhenoAge, GrimAgeV1, GrimAgeV2, YingCausAge, HorvathV1, Lin, Hannum, and HorvathV2; and (2) YingDamAge, DunedinPoAm38, Zhang10, and DunedinPACE. The DNAmTL telomere length clock and YingAdaptAge do not cluster with other clocks, suggesting their unique biological underpinnings.

Lastly, we compared the predictive power of aging biomarkers for healthspan and lifespan (Figure 4h). In general, we observed a significant positive correlation using an inverse-variance-weighted approach (weighted correlation coefficient = 0.58, P = 0.012). DunedinPoAm38, GrimAgeV2, GrimAgeV1, and Zhang_10 demonstrated strong associations with healthspan and lifespan, indicating their potential as comprehensive aging biomarkers. These findings highlight the utility of Biolearn in facilitating the systematic evaluation of aging biomarkers for predicting various health outcomes. The results provide insights into the comparative performance of these biomarkers and their potential applications in clinical settings and aging research.

### Transcriptomic and Clinical Biomarkers

To demonstrate Biolearn’s multi-omic capabilities, we also evaluated the performance of transcriptomic and phenotypic aging biomarkers. We include four transcriptomic age predictors: tAge.Peters ^30^, tAge.Multispecies.Blood, tAge.Human.Multi-tissue, and tAge.Human.Blood ^31,32^. We applied these predictors to the JenAge RNA-Seq dataset (Jena Centre for Systems Biology of Ageing, Illumina TruSeq 2.0, Whole Blood, N = 62) ^33^. The tAge predictor showed a strong correlation with chronological age, with Pearson’s R ranging from 0.68 (Peters) to 0.90 (Human. Multi-tissue) (Figure 5a), highlighting their potential as robust aging biomarkers. All these predictors are implemented in Biolearn with simple and easy-to-use functions (Figure 5b).

**Figure 5.**
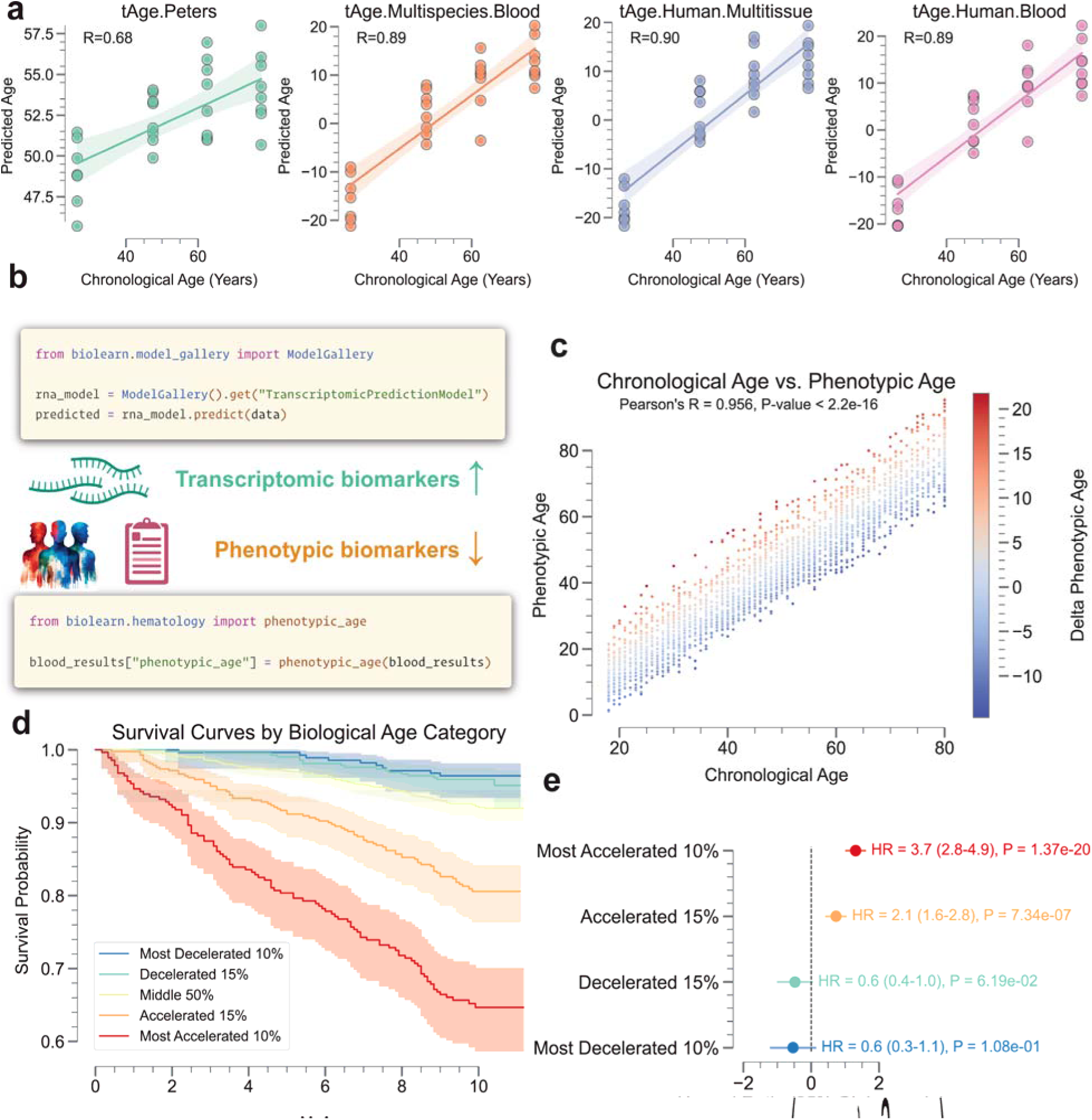
Transcriptomic and clinical biomarkers. **a.** Scatter plots of chronological age vs. predicted age for various transcriptomic clocks (tAge) on JenAge RNA-Seq dataset (Jena Centre for Systems Biology of Ageing, Illumina TruSeq 2.0, Whole Blood, N=62, GSE103232 & GSE75337). Pearson’s R values are shown at the top of the plot. **b.** Overview of Biolearn’s transcriptomic and phenotypic biomarker functionalities. The code snippet shows that transcriptomic age predictors like Peters (tAge) can be calculated with a few lines of code using the Biolearn library. **c.** Scatter plot showing the correlation between Phenotypic Age prediction (X-axis) and chronological age (Y-axis). The dots are colored by AgeDev. **d.** Survival analysis of the NHANES 2010 dataset (N = 2877), stratified by biological age discrepancies (marked by different colors) based on Phenotypic Age. The survival curves demonstrate that individuals with accelerated biological aging (red) have a lower survival probability compared to those with decelerated aging (blue). The shaded areas in c represent the standard error of the survival estimates. **e.** Forest plot displaying the hazard ratios (HR) and 95% confidence intervals (CI) for the association between biological age discrepancies based on the Phenotypic Age metrics and allcause mortality.

Next, we investigated the performance of the Phenotypic Age predictor ^11^, a blood-test-based biomarker, using the NHANES 2010 dataset. We calculated the Phenotypic Age for each individual using Biolearn’s implementation of the predictor and compared it to chronological age (Figure 5c). We found a strong linear relationship between Phenotypic Age and chronological age (Pearson’s R = 0.96, P < 2.2e-16), indicating the predictor’s ability to capture age-related changes in clinical biomarkers.

To assess the predictive power of Phenotypic Age on mortality risk, we performed a survival analysis on the NHANES 2010 dataset. Individuals were stratified into five groups based on their AgeDev (Figure 5d,e). We found that individuals with higher AgeDev had a significantly higher mortality risk (HR = 1.58, 95% CI: 1.31-1.91, P = 1.08e-06), while those with lower AgeDev had a lower risk (HR = 0.60, 95% CI: 0.49-0.74, P = 6.19e-07). These results underscore the predictive power of Phenotypic Age in assessing mortality risk and demonstrate Biolearn’s capability to facilitate such analyses.

To investigate the association between aging and cell type proportions, we performed deconvolution analysis on the NAS cohort blood samples. This analysis revealed distinct changes for different cell types over the life course (Extended Fig. 1). Neutrophil and natural killer cell proportions showed significant positive correlations with age (R=0.09, P=5.64e-04 and R=0.14, P=1.03e-07, respectively). In contrast, B cell, CD4 T cell, and CD8 T cell proportions exhibited significant negative correlations with age (R=-0.09, P=2.45e-04; R=-0.15, P=6.07e-09; and R= - 0.05, P=4.01e-02, respectively). Monocyte proportion did not show a significant correlation with age (R=0.02, P=4.07e-01).

We assessed the predictive power of cell type proportions on various health outcomes in the NAS cohort (Extended Fig. 2). For healthspan, natural killer cell proportion was a significant protective factor (HR=0.93, 95% CI: 0.88-0.99, P=1.55e-02), while CD8 T cell proportion was a slight risk factor (HR=1.06, 95% CI: 1.00-1.12, P=4.65e-02). Similarly, for lifespan, natural killer cell proportion was a significant protective factor (HR=0.91, 95% CI: 0.85-0.98, P=1.03e-02), and CD8 T cell proportion was a significant risk factor (HR=1.07, 95% CI: 1.01-1.14, P=2.59e-02). Natural killer cell proportion was also a significant protective factor for dementia (HR=0.87, 95% CI: 0.77-0.98, P=2.03e-02). For stroke risk, neutrophil proportion was a significant risk factor (HR=1.33, 95% CI: 1.11-1.60, P=2.18e-03), while natural killer cell proportion was a significant protective factor (HR=0.70, 95% CI: 0.55-0.89, P=4.21e-03). Similarly, for surgical events, neutrophil proportion was a significant risk factor (HR=1.16, 95% CI: 1.05-1.28, P=3.99e-03), and natural killer cell proportion was a significant protective factor (HR=0.89, 95% CI: 0.80-0.99, P=2.58e-02). No significant associations were found between cell type proportions and cancer risk.

The integration of transcriptomic and phenotypic biomarkers in Biolearn enables researchers to investigate aging processes from different biological perspectives. The strong performance of the tAge predictor and Phenotypic Age in their respective datasets showcases the potential of multiomic approaches in uncovering the complex mechanisms underlying aging. By leveraging Biolearn’s comprehensive framework, researchers can gain valuable insights into the interplay between different biological layers and their contributions to the aging process, ultimately facilitating the development of targeted interventions and personalized aging management strategies.

## Discussion

Among the most significant challenges in aging biomarker research is cross-population validation of proposed biomarkers ^34^. To take steps to address this need and provide an open-source tool for validation efforts across the field, we built Biolearn, an open-source library that provides a unified framework for the curation, harmonization, and systematic evaluation of aging biomarkers across diverse datasets. By leveraging Biolearn, we conducted a comprehensive benchmarking analysis of various epigenetic aging clocks, transcriptomic predictors, and phenotypic biomarkers, demonstrating their performance, robustness, and generalizability in different biological contexts and populations.

Systematic evaluation of epigenetic aging clocks across multiple datasets revealed that the Horvath skin and blood clock, Hannum clock, Horvath multi-tissue clock and Ying CausAge clock exhibited the highest overall accuracy in predicting chronological age. Notably, the predictive power of epigenetic clocks on mortality risk, after adjusting for age and sex, was independent of their ability to predict chronological age, highlighting the importance of interpreting AgeDev with caution and underscoring the need for further investigation into the biological mechanisms captured by these clocks. Evaluation of aging biomarkers in predicting various clinical outcomes in the NAS cohort revealed considerable heterogeneity, with different biomarkers showing strengths in predicting specific outcomes. For example, DunedinPACE was the strongest predictor for healthspan, dementia, and stroke, while PhenoAge and the Horvath skin and blood clock were the strongest predictors for surgery and cancer, respectively. These findings underscore the importance of selecting appropriate biomarkers based on the specific clinical outcomes of interest and suggest the potential for developing targeted interventions tailored to individual aging trajectories.

Integration of transcriptomic and phenotypic biomarkers in Biolearn enables a multi-omic approach to investigating the aging process. The strong performance of the tAge predictor and Phenotypic Age in their respective datasets highlights the potential of combining information from different biological layers to gain a more comprehensive understanding of the aging process. By leveraging Biolearn’s unified framework, researchers can explore the interplay between various omic modalities and uncover novel insights into the complex mechanisms underlying aging.

With Biolearn, we also harmonized and evaluated several well-established aging clocks, providing the opportunity for these biomarkers to be refined and potentially for new ones to be developed. The modular design of Biolearn encourages the addition of new models and datasets, making it a living library that will grow in tandem with the field itself. By centralizing resources and knowledge, Biolearn considerably reduces redundancy and accelerates biomarker development and validation efforts ^5,9^.

Our approach emphasizes transparency and reproducibility, core tenets of open science. By making Biolearn publicly available and maintaining detailed documentation and development guidelines, we established an ecosystem that supports open collaboration and knowledge sharing. This open-science framework ensures that findings and tools can be widely accessed, providing equitable opportunities for researchers globally to contribute to and benefit from the collective advances in aging research. Moreover, our hope is that the open-access nature of Biolearn will promote cross-fertilization between aging researchers and scientists currently outside the field, incentivizing the development of novel and innovative biomarker models and validation approaches.

Previous efforts to harmonize biomarkers of aging, notably methylCIPHER and BioAge ^35,36^, have been limited in scope, focusing on methylation or blood-based biomarkers only. Furthermore, using R packages is somewhat limiting. Biolearn supports biomarkers based on multiple different biological data modalities and is written in Python, which has a broader reach. In comparison to PyAging ^37^, a preliminary contemporaneous Python biomarker library, Biolearn is focused on ease of use and reproducibility through automated testing against reference data. Biolearn also offers several distinct advantages over existing biomarker libraries: it supports the easy loading of a larger and more diverse set of data, enabling researchers to work with a wide range of datasets and explore the performance of biomarkers across various populations and biological contexts; it includes models that are not directly used for age prediction but may be relevant to health and lifespan, providing a more comprehensive toolkit for aging research; and it includes tooling for exploring and understanding your data, such as quality reports, which facilitate data preprocessing and ensure the reliability of the results.

While Biolearn represents a significant advance for the field, some limitations remain. Currently, the library is tailored to biomarkers derived from biological samples, predominantly DNA methylation data. Moving forward, the scope of Biolearn will continue to expand to encompass diverse biological modalities—such as proteomics, metabolomics, and microbiomics—broadening its applicability ^7,13^. Moreover, integration with larger and more diverse population datasets will be vital in advancing cross-population validation efforts. As new datasets emerge, Biolearn will adapt to incorporate these resources, ensuring ongoing robustness and scalability ^38^. Finally, bioinformatics tools, including Biolearn, depend on a user base proficient in programming and data analysis. Efforts to make these tools more accessible to a wider audience, including those with limited computational expertise, will be crucial. This could involve the development of graphical user interfaces (GUIs) or web-based platforms to streamline the user experience.

We anticipate that Biolearn will become a key resource for the field and will transform many facets of aging biomarker studies. Our preliminary survival studies conducted using Biolearn demonstrate not only the power of this new platform but illuminate the real-world implications of validated biomarkers. Biolearn’s standardization and analysis capabilities stand to serve as pivotal tools for researchers seeking to bridge the gap between biomarker discovery and clinical implementation ^34^.

## Methods

### Overview of Biolearn Library

Biolearn is an open-source computational suite that facilitates the harmonization and analysis of biomarkers of aging (BoAs). It is written in Python and is readily accessible through the Python Package Index (PyPI). Biolearn is developed using modern software engineering practices, including automated testing to ensure correctness and adherence to software design principles that ensure the safe interchangeability of like components. The library is designed to be user-friendly while offering robust functionalities for researchers across various disciplines involved in aging studies. Additionally, the results of each model in Biolearn are portable and can be easily exported in formats such as CSV, allowing for seamless integration with other tools like R for further analysis and visualization.

### System Requirements and Installation

Biolearn requires Python version 3.10 or newer. It can be installed using the Python package manager, pip, with the command pip install biolearn. The successful installation of the library can be verified through the import test of Biolearn’s core classes. The library is cross-platform and is compatible with major operating systems, including Windows, MacOS, and Linux.

### Data Library and Model Gallery

Biolearn incorporates a data library capable of loading and structuring datasets from a multitude of public sources like Gene Expression Omnibus (GEO), National Health and Nutrition Examination Survey (NHANES), and Framingham Heart Study. The model gallery within Biolearn holds reference implementations for various aging clocks and biomarkers, presenting a unified interface for users to apply these models to their datasets. All models were verified to be correct by comparing the outputs on a reference data set against their original implementations where available.

### Harmonization Process

We used Biolearn to harmonize several aging clocks. Clock definitions were standardized, specifying the name, publication year, applicable species, target tissue, and the biological aspect they predict (e.g., age, mortality risk). We provided sources for both the original publications and the coefficients necessary for clock applications. Coherence across biological modalities and datasets was assured through Biolearn’s systematic approach to data preprocessing, normalization, and imputation procedures.

### Integration with Public Datasets

Biolearn’s ability to interface seamlessly with public datasets was tested by integrating and formatting data from GEO and NHANES. Preprocessing pipelines were developed to convert raw data into a harmonized format suitable for subsequent analysis. Particular attention was given to metadata structures, variable normalization, and missing data treatment, ensuring consistent input formats required by the aging models.

### Cell Type Deconvolution

Biolearn’s deconvolution function estimates cell type proportions within a sample from bulk methylation data. It operates on the principle that bulk methylation is a composite of methylation profiles from various cell types, proportionate to their presence (B = X * P). Here, ‘B’ represents bulk methylation, ‘X’ is the matrix of cell-type-specific methylation profiles, and ‘P’ is the vector of cell-type proportions. With known ‘B’ and estimated ‘X,’ ‘P’ is computable through constrained quadratic programming, ensuring proportions remain within biological plausibility— between zero and one and summing to one ^26^.

We incorporated two deconvolution variants tailored for blood methylation analysis: DeconvoluteBlood450K for the 450K platform and DeconvoluteBloodEPIC for the EPIC platform, addressing platform-specific biases ^27^. The methylation matrix for each mode derives from the corresponding platform, encapsulating six major blood cell types—neutrophils, monocytes, NK cells, B cells, CD4+ T cells, and CD8+ T cells. Selection of CpG sites for deconvolution relied on identifying the 50 most distinctively hyper- and hypo-methylated sites per cell type, prioritized by the significance of their differential methylation. This yielded 600 reference CpG sites per deconvolution mode ^28,29^.

### Statistical Analysis

All statistical analyses were performed using tools embedded within the Biolearn library or through integration with renowned Python statistics libraries such as statsmodels and seaborn for visualization. The robustness and reproducibility of the analysis were ensured through the use of randomized cross-validation techniques for model assessment and bootstrapping methods for estimating confidence intervals where applicable. Survival analyses were conducted using the Cox Proportional Hazards model, adjusting for age and other relevant covariates. The performance of aging clocks in predicting chronological age and mortality risk was evaluated using metrics such as R^2^, hazard ratios, and p-values.

## Acknowledgments

We thank all Biomarkers of Aging Consortium members for their valuable feedback and suggestions. We thank S. Horvath and A. Lu for sharing the GrimAgeV1 and V2. This work was inspired by methylCIPHER, an R package for DNA methylation clocks ^36^. Supported by grants from the National Institute on Aging, Hevolution Foundation, Methuselah Foundation, and VoLo Foundation.

## Notes

https://bio-learn.github.io/

### Competing Interest Statement

The authors have declared no competing interest.

### Summary of Updates

Major update of the analysis. Removed MGB result for the competition

